# A ubiquitin-independent proteasome pathway controls the CARD8 inflammasome

**DOI:** 10.1101/2022.03.22.485305

**Authors:** Jeffrey C. Hsiao, Atara R. Neugroschl, Ashley J. Chui, Cornelius Y. Taabuzing, Andrew R. Griswold, Qinghui Wang, Hsin-Che Huang, Elizabeth L. Orth-He, Daniel P. Ball, Giorgos Hiotis, Daniel A. Bachovchin

## Abstract

CARD8 is a pattern-recognition receptor that forms a caspase-1-activating inflammasome. CARD8 undergoes autoproteolysis, generating an N-terminal (NT) fragment with a disordered region and a ZU5 domain and a C-terminal (CT) fragment with UPA and CARD domains. DPP8 and DPP9 (DPP8/9) inhibitors, including Val-boroPro (VbP), accelerate the degradation of the NT fragment via a poorly characterized proteasome-mediated pathway, thereby releasing the inflammatory CT fragment from autoinhibition. Here, we show that the core 20S proteasome, which degrades disordered and misfolded proteins independent of ubiquitin, controls CARD8 activation. In unstressed cells, the 20S proteasome degrades just the NT disordered region, leaving behind the folded ZU5, UPA, and CARD domains to act as an inhibitor of inflammasome assembly. In VbP-stressed cells, the 20S proteasome degrades the entire NT fragment, perhaps due to ZU5 domain unfolding, freeing the CT fragment from autoinhibition. Overall, this work shows that CARD8 NT’s susceptibility to 20S proteasome-mediated degradation controls inflammasome activation.

## Introduction

Several intracellular danger-associated signals induce the assembly of multiprotein complexes called inflammasomes (1,2). The typical process of inflammasome formation involves a pattern recognition receptor (PRR) protein detecting a specific danger signal, self-oligomerizing, and then recruiting (directly or indirectly via the adapter protein ASC) the cysteine protease caspase-1 (CASP1). CASP1 undergoes proximity-induced autoproteolysis on this platform, generating an active enzyme that cleaves and activates gasdermin D (GSDMD) and, in most cases, interleukin-1β (IL-1β) and IL-18. The N-terminal fragment of cleaved GSDMD (GSDMD^p30^) forms pores in the cell membrane, releasing the activated cytokines and triggering pyroptotic cell death.

CARD8 is a human PRR that forms an inflammasome (3). CARD8 has an N-terminal unstructured region consisting of ∼160 amino acids followed by a function-to-find domain (FIIND) and a caspase activation and recruitment domain (CARD) (**Fig. 1A**). The FIIND undergoes autoproteolysis between its ZU5 (found in ZO-1 and UNC5) and UPA (conserved in UNC5, PIDD, and ankyrins) subdomains, creating N-terminal (NT) and C-terminal (CT) fragments that remain non-covalently associated (4). The proteasome-mediated degradation of the NT fragment releases the CT fragment from autoinhibition, but the CT fragment is then captured and restrained as part of a ternary complex with one copy of full-length (FL) CARD8 and one copy of dipeptidyl peptidase 8 or 9 (DPP8/9) (5). Stimuli that accelerate CARD8^NT^ degradation and/or disrupt the DPP8/9-CARD8 complex enable the CARD8^CT^ to overcome these repressive mechanisms and to self-oligomerize, recruit CASP1, and trigger pyroptosis.

**Figure 1.**
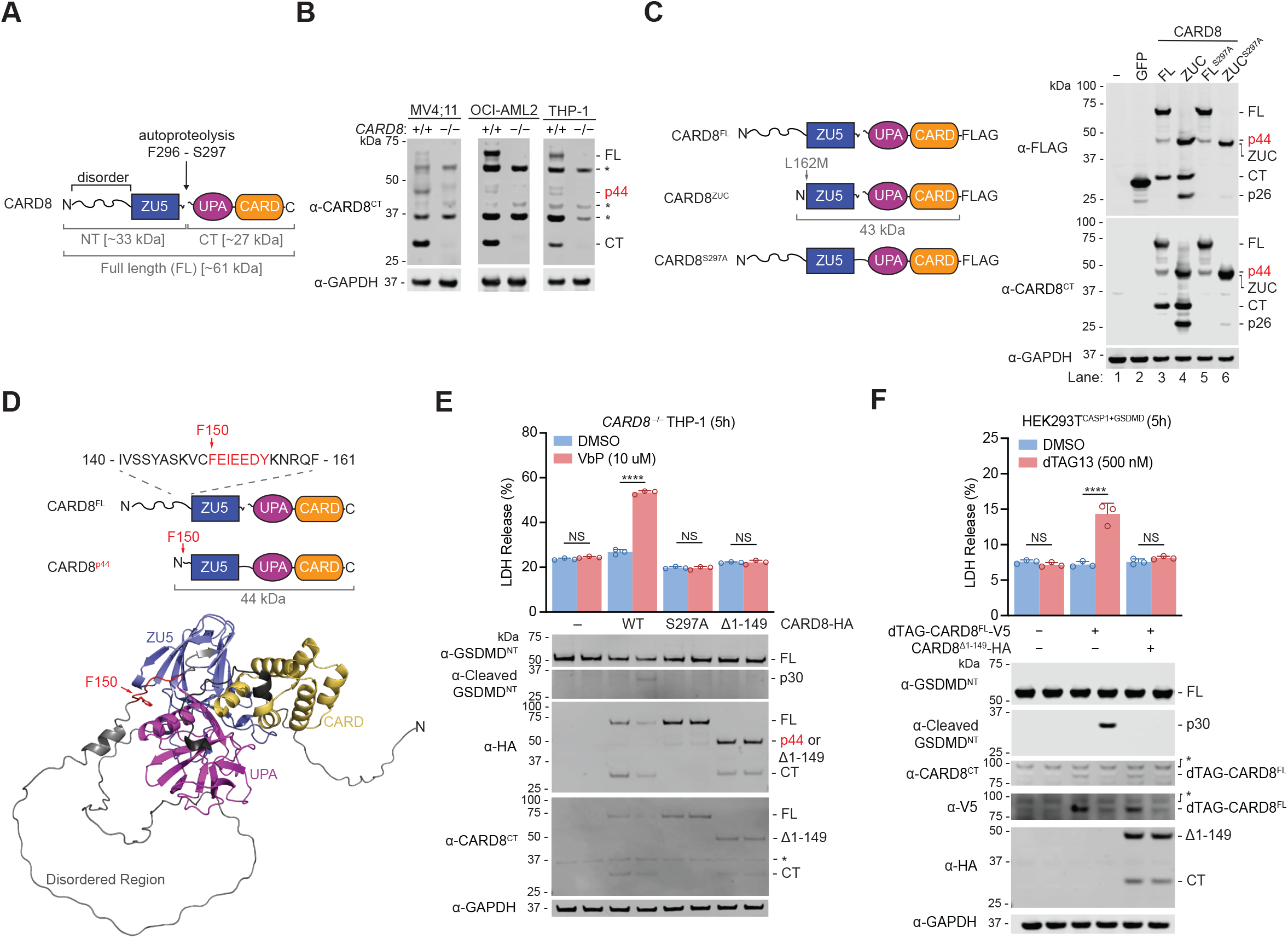
CARD8’s disordered region is removed in cells. (**A**) Domain organization of CARD8. CARD8 undergoes autoproteolysis between the ZU5 and UPA subdomains. The size in kilodaltons (kDa) of each fragment is indicated. (**B**) Lysates of the indicated cell lines were analyzed by immunoblotting. (**C**) HEK 239T cells were transfected with plasmids encoding the indicated FLAG-tagged constructs (left). Lysates were analyzed by immunoblotting (right). (**D**) Edman degradation analysis revealed residues F150 to Y156 (colored red) as the N-terminus CARD8^p44^. F150 is shown on the CARD8 structure predicted by Alpha-Fold (23). (**D**) *CARD8*^*−/−*^ THP-1 cells stably expressing CARD8 WT, S297A, or β1-149 were treated with compounds for 5 hours before LDH release and immunoblotting analyses. (**F**) HEK 293T^CASP1+GSDMD^ cells were transfected with plasmids encoding the indicated plasmids. After 24 h, cells were treated with dTAG-13 for 5 h before LDH release and immunoblot analyses. Data in (**E**) (n=3) and (**F**) (n=3) are means ± standard deviation (SD) of replicates. **** *p* < 0.0001, by Student’s two-sided *t*-test. NS, not significant. All data, including immunoblots, are representative of three or more independent experiments.

Two distinct danger signals have been reported to accelerate the proteasome-mediated degradation of the CARD8^NT^ fragment. First, HIV-1 protease directly cleaves within the NT region of the CARD8^FL^ protein, generating an unstable neo-N-terminus that is rapidly degraded by the N-end rule proteasome pathway (6). Second, DPP8/9 inhibitors, including Val-boroPro (VbP), accelerate the degradation of many disordered and misfolded proteins, including the CARD8^NT^(7,8). Notably, DPP8/9 inhibitors also destabilize the repressive DPP8/9-CARD8 ternary complex (5), and thereby activate the CARD8 inflammasome via two separate mechanisms.

The molecular details of the homeostatic and DPP8/9-inhibition induced CARD8 degradation pathways have not been established. Intracellular proteins are often degraded by the ubiquitin-proteasome system (UPS), which involves the covalent attachment of ubiquitin to lysine residues on target proteins that mediate their recruitment to the 26S proteasome. The 26S proteasome consists of the proteolytic core 20S subunit capped at one or both ends by 19S regulatory complexes (9). The 19S regulatory particles recognize, deubiquitinate, and unfold target proteins, enabling their translocation into the 20S core particle for hydrolysis. In a preliminary attempt to identify sites of ubiquitination on CARD8, we previously mutated all 10 lysines within the NT fragment of CARD8^FL^ to arginines (CARD8^FL^ K10R). We found that CARD8^FL^ K10R was largely, but not completely, insensitive to VbP in a reconstituted HEK 293T cell system expressing CASP1 and GSDMD (HEK 239T^CASP1+GSDMD^ cells), suggesting that ubiquitination of the NT fragment might be important for VbP-induced degradation (8). However, the CARD8^FL^ K10R protein expressed at lower levels and underwent less autoproteolysis than the wild-type (WT) CARD8^FL^ protein, and these deficiencies could also account for its reduced pyroptotic activity.

Here, we further investigated the molecular details of CARD8 degradation. We found that the core 20S proteasome, which degrades misfolded proteins independent of the 19S regulatory complex and ubiquitination, regulates CARD8 activation. In unstressed cells, the 20S proteasome removes the disordered region of CARD8, leaving behind the folded ZU5-UPA-CARD (ZUC) domains. This protein fragment cannot form an inflammasome but can still sequester CT fragments in the DPP8/9 ternary complex and thereby act as an inflammasome inhibitor. In VbP-stressed cells, the 20S proteasome degrades CARD8’s entire NT fragment, including the ZU5 domain, releasing the inflammatory CT fragment from autoinhibition. Collectively, these findings suggest that the propensity of the ZU5 domain to enter the 20S proteasome is a critical regulatory step that governs the activation of the CARD8 inflammasome.

## Results

### CARD8’s disordered region is removed in cells

Before further studying VbP-induced CARD8^NT^ degradation, we first wanted to investigate how CARD8 is processed in unstressed cells. Intriguingly, we and others have consistently observed that endogenous CARD8 in human monocytes appears as three distinct bands at 61, 44, and 27 kDa in immunoblots using antibodies targeting CARD8^CT^ (**Fig. 1B**) (3,4,10). Only a fraction of the CARD8^FL^ typically undergoes autoproteolysis (4,11,12), and the bands at 61 kDa and 27 kDa correspond to CARD8^FL^ and CARD8^CT^, respectively. The identity of the third band at 44 kDa, which we call CARD8^p44^, however, is unknown. CARD8^p44^ is unlikely to be a splicing isoform, as the transfection of the cDNA encoding the canonical 61 kDa CARD8 isoform (i.e., isoform 5) in HEK 293T cells also generated CARD8^p44^ (**Fig. 1C**, lane 3). Notably, ectopic expression of the autoproteolysis-defective S297A mutant CARD8^FL^ protein (CARD8^FL^ S297A) similarly generated CARD^p44^ (**Fig. 1C**, lane 5), demonstrating that formation of this species does not depend on autoproteolysis. We predicted that CARD8^p44^ corresponded to CARD8^FL^ protein that was N-terminally truncated prior to its ZU5 domain because it migrated slightly more slowly than the isolated ZU5-UPA-CARD domains (CARD8^ZUC^) (**Fig. 1C**, lanes 4 and 6) and was not detected with antibodies targeting CARD8^NT^ (**Fig. S1A**). Indeed, Edman degradation analysis revealed that the N-terminal residue of CARD8^p44^ was F150 (**Fig. 1D** and **Fig. S1B**). Thus, a fraction of CARD8^FL^ is proteolytically cleaved twelve residues before the start of the ZU5 domain to generate CARD8^p44^. It should be noted that the isolated CARD8^ZUC^ also appears to further be processed into a p26 fragment, but the identity and function of this band was not studied further here (**Fig. 1C**, lane 4).

We previously evaluated the abilities of several N-terminally truncated CARD8 proteins to mediate VbP-induced pyroptosis in HEK 239T^CASP1+GSDMD^ cells (8). In this analysis, we discovered a CARD8 construct starting at K147, but not F150, was capable of mediating pyroptosis, suggesting that CARD8^p44^ is likely incapable of forming an inflammasome. We next wanted to confirm this result in THP-1 cells, which endogenously express the CARD8 inflammasome pathway and are therefore more physiologically relevant (3). To do this, we ectopically expressed CARD8^FL^ WT, CARD8^FL^ S297A, or CARD8 lacking residues 1-149 (CARD8^Δ1-149^) in *CARD8*^*−/−*^THP-1 cells before treating the cells with DMSO or VbP (**Fig. 1E**). As expected, VbP induced pyroptosis in THP-1 cells expressing CARD8^FL^ WT, but not CARD8^FL^ S297A or CARD8^Δ1-149^. The aminopeptidase inhibitor bestatin methyl ester (MeBs) synergizes with VbP to induce more pyroptosis (7), but the combination of VbP and MeBs still failed to activate CARD8^Δ1-149^ (**Fig. S1C**). Collectively, these data show that CARD8^p44^ cannot form an inflammasome.

The CARD8^ZUC^ can occupy the CARD8^FL^ position in the CARD8-DPP8/9 ternary complex, and thus can capture and repress a freed CARD8^CT^ fragment (5). As such, we predicted that CARD8^p44^, which is only slightly longer than CARD8^ZUC^, might function as an inhibitor of inflammasome formation. To test this idea, we expressed CARD8^FL^ with an N-terminal degradation tag (dTAG-CARD8^FL^) in HEK 239T^CASP1+GSDMD^ cells (**Fig. 1F**) (5,13). The small molecule dTAG-13 triggers the rapid degradation of proteins fused to dTAGs, and therefore treatment of these cells with dTAG-13 induced the release of free CARD8^CT^ and pyroptosis (**Fig. 1F**). Consistent with our hypothesis, the co-expression of CARD8^Δ1-149^ in these cells abolished pyroptosis without impacting dTAG-13-induced dTAG-CARD8^FL^ degradation (**Fig. 1F**). Overall, these results suggest than an endogenous protease removes CARD8’s disordered region and generates an inhibitory form of CARD8 that blocks inflammasome activation.

### The proteasome generates CARD8^p44^

We next wanted to determine the sequence and structural requirements for the proteolysis of CARD8^FL^ into CARD8^p44^. We previously created the chimeric protein MTMR1^M1-Q94^-CARD8^ZUC^, in which the disordered region of CARD8 was replaced by the disordered region (residues M1 to Q94) of MTMR1 (8). Intriguingly, the expression of this chimeric protein in HEK 293T cells still generated a p44 fragment (**Fig. 2A**, lane 3). The disordered regions of MTMR1 and CARD8 do not share any homology (**Fig. S2A**), indicating that the unknown protease does not recognize and cleave a specific amino acid sequence. To determine if the CARD8^ZUC^ specifically directs the cleavage, we appended the disordered region of CARD8 on the N-terminus of GFP. Interestingly, we found that that the disordered region was similarly removed to generate a protein species with a molecular weight close to GFP’s (**Fig. S2B**). In addition, we found that appending a well-folded GFP tag to the N-terminus of CARD8^FL^ or CARD8^Δ1-130^ did not interfere with CARD8^p44^ generation (**Fig. 2A**, lanes 4 and 5), showing that the unknown protease has endo-peptidase activity (i.e., it does not only remove disordered regions from protein N-termini). In contrast, replacement of the entire CARD8 disordered region with GFP (GFP-CARD8^ZUC^) abolished CARD8^p44^ generation (**Fig. 2A**, lane 6). Notably, an antibody targeting GFP detected bands corresponding to GFP itself for the GFP-CARD8 chimeras in which some disorder is present (i.e., GFP CARD8^FL^ and GFP CARD8^Δ1-130^), but not from the GFP-CARD8^ZUC^ protein (**Fig. S2C**, lanes 2 to 4). Overall, these data indicate that the CARD8-cleaving protease removes disordered regions in a sequence independent manner, including those between two well-folded domains.

**Figure 2.**
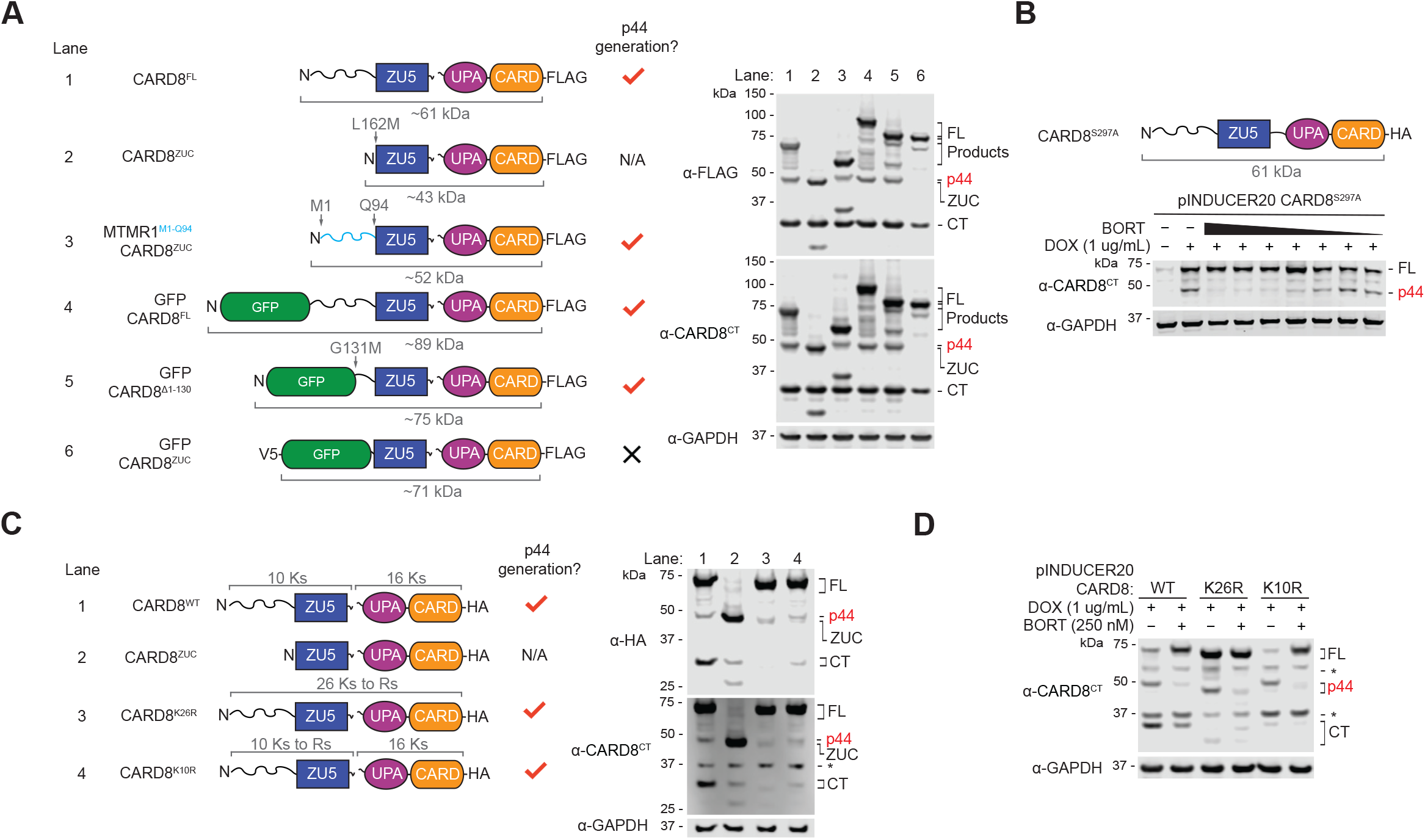
The proteasome generates CARD8^p44^ in cells. (**A**) HEK 239T cells were transiently transfected with plasmids encoding the indicated FLAG-tagged constructs (left). Lysates were analyzed by immunoblotting (right). The lane and p44 generation for each construct are shown. (**B**) A tetracycline-on (tet-on) plasmid encoding CARD8^FL^ S297A was transfected into HEK 293T cells. After 20 h, cells were treated with doxycycline (DOX) and/or Bortezomib (BORT). BORT was used in a 7-point, 3-fold dilution series from 1 uM to 1.37 nM. (**C**) HEK 239T cells were transiently transfected with plasmids encoding the indicated HA-tagged constructs (left). Lysates were analyzed by immunoblotting (right). The lane and p44 generation for each construct are shown. (**D**) HEK 293T cells were transiently transfected with tet-on plasmids encoding the indicated CARD8 proteins. After 20 h, cells were treated with doxycycline (DOX) and/or Bortezomib (BORT). Lysates were harvested after 24 h and analyzed with the indicated antibodies. Immunoblots are representative of three or more independent experiments.

Intriguingly, the proteolytic 20S core of the proteasome, which often functions independently of the 19S cap in cells, degrades unstructured polypeptides regardless of amino acid sequence, including those between structured domains (14-16). As such, we hypothesized that the 20S proteasome was the unknown protease generating CARD8^p44^. To test this idea, we transiently transfected HEK 293T cells with a doxycycline (DOX)-inducible construct encoding autoproteolysis-defective CARD8^FL^ S297A, treated cells with DOX and with increasing concentrations of the proteasome inhibitor bortezomib, and assayed for newly formed CARD8^p44^ by immunoblotting (**Fig. 2B**). We observed that bortezomib blocked the generation of CARD8^p44^, strongly indicating that the proteasome was indeed responsible for this cleavage.

The 26S proteasome typically requires the covalent attachment of ubiquitin to substrate proteins preceding their degradation, whereas the 20S proteasome directly degrades misfolded proteins without ubiquitination (14-16). To determine whether CARD8^p44^ formation involves lysine ubiquitination, we mutated all lysines to arginines within the NT fragment (CARD8^FL^ K10R) or throughout the entire protein (CARD8^FL^ K26R). It should be noted that the CARD8^FL^ K26R protein is entirely devoid of lysines, including its C-terminal linker and hemagglutinin (HA)-tag. We observed that both mutant proteins still generated CARD8^p44^ fragments in HEK 293T cells, and that bortezomib still attenuated the formation of these products (**Fig. 2C,D**). These results suggest that the 20S proteasome removes the disordered region of CARD8 through a ubiquitin-independent mechanism.

### The 20S proteasome generates CARD8^p44^ in vitro

As proteasome inhibition might indirectly block CARD8^p44^ formation in cells, we next sought to confirm that the purified 20S proteasome directly removes CARD8’s disordered region to generate CARD8^p44^ *in vitro*. We therefore purified a C-terminally FLAG-tagged CARD8^FL^ protein from HEK 293T cells using anti-FLAG beads, and then incubated this protein with purified 20S proteasomes. We found that the 20S proteasome robustly degraded CARD8^FL^ into CARD8^p44^, but that the presumably well-folded CARD8^p44^ and CARD8^CT^ products were largely protected from degradation (**Fig. 3A,B**). It should be emphasized that the purified 20S proteasome generated a p44 band precisely the same size as the one present from constitutive processing by endogenous proteasomes, strongly indicating that the 20S proteasome generates CARD8^p44^ in cells. As expected, the proteasome inhibitors bortezomib and MG-132 slowed the *in vitro* generation of CARD8^p44^ (**Fig. S3**). Like CARD8^p44^ and CARD8^CT^, we found that the CARD8^ZUC^ was resistant to 20S proteasome-mediated degradation for at least 4 h (**Fig. 3B**).

**Figure 3.**
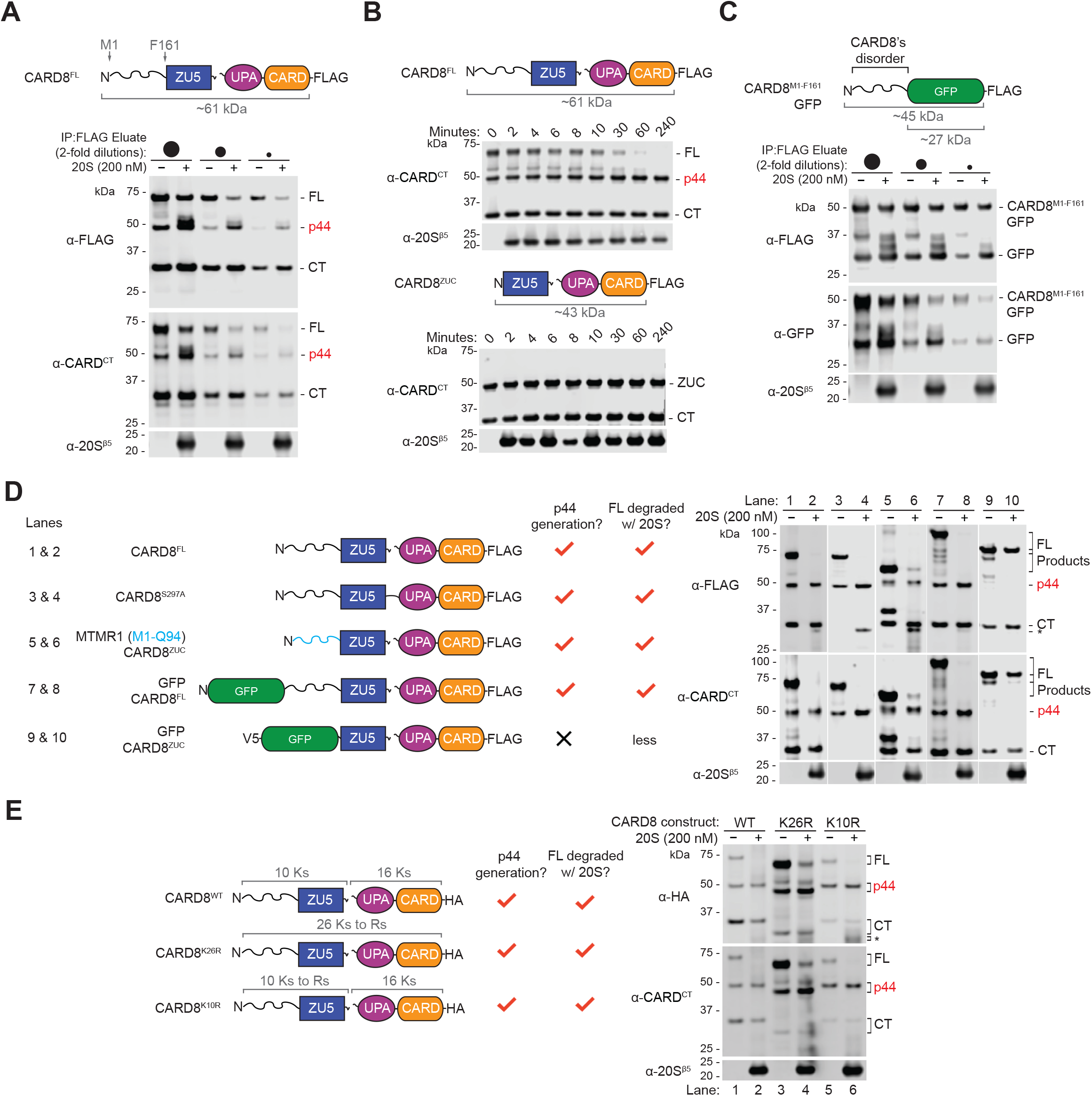
The 20S proteasome generates CARD8^p44^ *in vitro*. (**A**) Purified CARD8^FL^ protein were incubated at varying dilutions with or without purified 20S proteasomes for 4 h. Reactions were quenched with 2X loading dye prior to immunoblotting analysis. (**B**) Purified CARD8^FL^ and CARD8^ZUC^ (each 800 nM) were incubated with purified 20S proteasomes (100 nM). At indicated timepoints, aliquots were removed from the mixture, quenched with 2X loading dye, and analyzed by immunoblotting. (**C-E**) The indicated purified proteins were treated and analyzed as described in (**A**). Immunoblots are representative of three or more independent experiments.

As mentioned above, the 20S proteasome has previously been shown to degrade disordered sequences, but spare well-folded domains (14,15). To determine if the 20S proteasome could indeed process the chimeric proteins evaluated above, we purified several of these proteins from HEK 293T cells and similarly incubated them with purified 20S proteasomes. As expected, we found that the purified 20S proteasome efficiently removed the CARD8 disordered region (M1-F161) fused to the N-terminus of GFP (**Fig. 3C**), as well as the MTMR1 disordered region (M1-Q94) fused to the N-terminus of CARD8^ZUC^ (**Fig. 3D**, lane 6). Also as expected, the 20S proteasome exhibited endoproteolytic activity, generating CARD8^p44^ even when a GFP-tag was appended to the N-terminus of CARD8^FL^ (**Fig. 3D**, lane 8). In stark contrast, GFP-CARD8^ZUC^, a chimeric protein that lacks the internal disordered region, was resistant to 20S proteasome-mediated degradation (**Fig. 3D**, lane 10). Lastly, 20S proteasomes processed the CARD8^FL^ K10R and K26R proteins into p44 fragments, in agreement with the known capability of the 20S to degrade substrates without lysines or ubiquitination.

### VbP activates CARD8 lacking NT lysines

Interestingly, the 20S proteasome did not always appear to generate substantially more of CARD8^p44^ from CARD8^FL^ *in vitro* (e.g., **Fig. 3D**, lanes 1 and 2**; Fig. 3E**, lanes 1 and 2), suggesting that degradation may proceed through the ZU5 domain in some cases. Moreover, VbP does not induce visible ubiquitination of CARD8 by immunoblotting (**Fig. S4A**) (3,5,7,8,10). We therefore wanted to determine if the 20S proteasome also mediates VbP-induced degradation of the entire CARD8^NT^ fragment, the key process that releases the pyroptotic CARD8^CT^ fragment from autoinhibition. To further explore this idea, we more closely investigated the ability of CARD8^FL^ K10R to stimulate pyroptosis. As mentioned above, we previously found that CARD8^FL^ K10R was largely defective in mediating VbP-induced pyroptosis in the reconstituted HEK 293T^CASP1+GSDMD^ system (8), but this inactivity was perhaps due to compromised autoproteolysis or expression. Here, we instead stably expressed CARD8^FL^ K10R in the more physiologically relevant *CARD8*^*−/−*^ THP-1 cells. We observed that VbP, and especially the combination of VbP and MeBs, induced pyroptosis, as evidenced by LDH release and GSDMD cleavage, in cells expressing CARD8^FL^ K10R (**Fig. 4A,B**). Moreover, we found that bortezomib (and caspase-1 inhibitor VX-765) abolished CARD8^FL^ K10R-dependent pyroptosis (**Fig. 4A**), showing that the lysine-free CARD8^NT^ fragment was indeed being degraded by the proteasome. As expected, neither the autoproteolysis defective CARD8^FL^ K10R/S297A nor the p44 fragment of CARD8^FL^ K10R (CARD8^Δ1-149^ K3R) formed inflammasomes (**Fig. 4A**).

**Figure 4.**
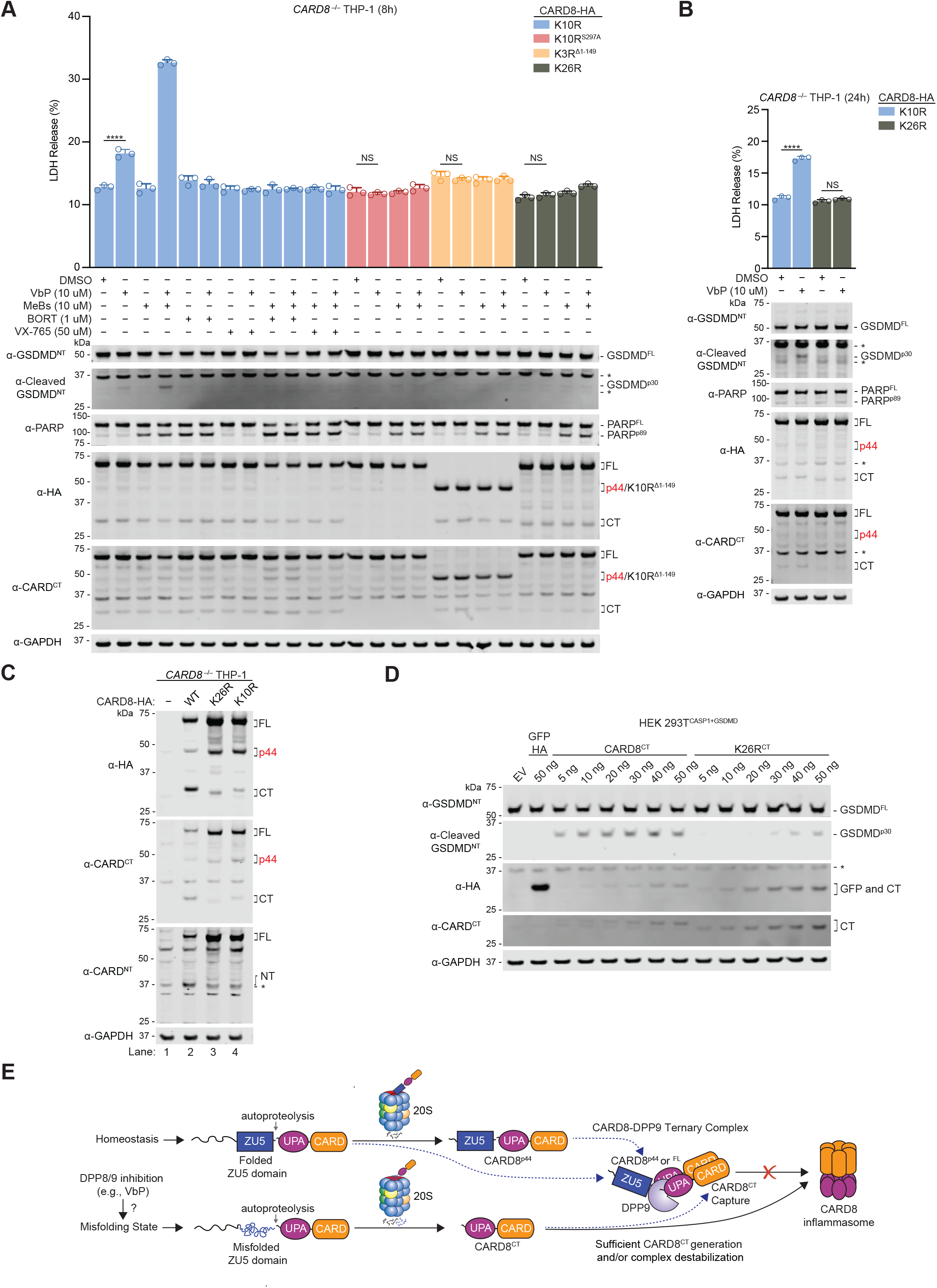
VbP activates CARD8^FL^ lacking NT lysines. (**A**) *CARD8*^*−/−*^ THP-1 stably expressing the indicated constructs were treated with the compounds for 8 hours before LDH release and immunoblotting analyses. PARP (poly ADP-ribose polymerase 1) cleavage to its p89 fragment was assessed as an indicator of apoptosis. (**B**) *CARD8*^*−/−*^ THP-1 cells stably expressing the indicated constructs were treated with the indicated compounds for 24 h before LDH release and immunoblotting analyses. Data in (**A**) and (**B**) are means ± standard deviation (SD) of three replicates. **** *p* < 0.0001 by Student’s two-sided *t*-test. NS, not significant. (**C**) Lysates of *CARD8*^*−/−*^ THP-1 cells stably expressing the indicated constructs were analyzed by immunoblotting. (**D**) HEK 293T^CASP1+GSDMD^ cells were transiently transfected with plasmids encoding the WT and K26R CT fragments and incubated for 16 h prior to immunoblotting analyses. All data, including immunoblots, are representative of three or more independent experiments. (**E**) Proposed model for the regulation of CARD8 by the 20S proteasome. In unstressed cells, the 20S proteasome removes the disordered region to generate CARD8^p44^, which acts as an inhibitor in the DPP8/9 ternary complex. In VbP-stressed cells, the 20S degrades the entire CARD8^NT^ fragment, possibly due to ZU5 domain misfolding, releasing CARD8^CT^ from autoinhibition. Increased degradation of the full NT fragment enables the CT fragment to overcome the ternary complex.

In theory, it remained possible that VbP induces the ubiquitination of the CT fragment of the auto-proteolyzed CARD8^FL^ K10R protein, and that this ubiquitination is sufficient to recruit CARD8 to the 26S proteasome and trigger its degradation in an N to C-terminus direction. To test this idea, we next investigated the ability of the lysine-free CARD8^FL^ K26R protein to mediate pyroptosis. We found that neither VbP nor the combination of VbP and MeBs induced pyroptosis in *CARD8*^*−/−*^ THP-1 cells ectopically expressing CARD8^FL^ K26R (**Fig. 4A,B**). However, this protein was likely inactive for at least two reasons. First, CARD8^FL^ K26R was severely defective in autoproteolytic activity (**Fig. 4C**, lanes 2 and 3**; Fig. S4B** lanes 3 and 6; densitometry ratio of CT:FL in CARD8^FL^ WT and K26R are 60% and 6%, respectively). Second, the isolated CT fragment of this construct (which contains sixteen lysines mutated to arginines) was impaired in its ability to oligomerize into a functional inflammasome, as transient transfection of considerably more plasmid encoding this mutant CT fragment relative to the WT CT fragment was required to induce GSDMD cleavage in HEK 239T^CASP1+GSDMD^ cells (**Fig. 4D**). Moreover, despite its other deficiencies, CARD8^FL^ K26R still retained binding to DPP9 and acted as an inflammasome inhibitor in the ternary complex (**Fig. S4C,D**). Collectively, these results suggest that the inability of CARD8^FL^ K26R to mediate pyroptosis is not necessarily due to a lack of ubiquitination sites. Because this lysine-free protein is non-functional for several reasons, it is unfortunately not possible to unequivocally demonstrate that VbP-induced NT degradation is independent of lysine ubiquitination.

We previously discovered that the NEDD8-activating enzyme inhibitor MLN4924 (17) blocks VbP-induced CARD8 activation (8). Cullin E3 ligases require neddylation for their activity, and we therefore speculated that a cullin E3 ligase might ubiquitinate CARD8. Notably, the activity of MLN4924 is the only evidence that suggests that CARD8’s degradation requires ubiquitination. Here, we found that MLN4924 blocks VbP-induced pyroptosis in *CARD8*^−/−^ THP-1 cells expressing CARD8^FL^ K10R (**Fig. S4E**). Therefore, we reasoned that MLN4924 treatment might inhibit CARD8 inflammasome activation by indirectly disrupting 20S proteasome activity, and not by directly abrogating CARD8 ubiquitination. Supporting this hypothesis, we found that MLN4924 reduced the formation of CARD8^p44^, showing that it likely indirectly interferes with 20S proteasome activity in cells (**Fig. S4F**). Overall, our discovery that CARD8^FL^ K10R is functional, coupled with the lack of VbP-induced ubiquitination, strongly indicates that the 20S proteasome mediates VbP-induced pyroptosis independent of the ubiquitin-proteasome system.

## Discussion

The VbP-induced proteasome pathway that degrades CARD8 has not yet been established. We previously speculated that VbP activates some unknown E3 ligase that specifically ubiquitinates the CARD8^NT^ fragment, thereby sending it to the 26S proteasome for destruction (8). However, no direct evidence showing that the ubiquitin-proteasome system is involved in this degradation process has emerged. Here, we show that the core 20S proteasome, which destroys unfolded or misfolded proteins independent of ubiquitin (14,15), controls CARD8 inflammasome activation.

Our proposed model for the regulation of CARD8 by the 20S proteasome is shown in **Figure 4E**. In unstressed cells during normal homeostasis, the CARD8 ZU5, UPA, and CARD domains fold properly, but the N-terminal disordered region remains unstructured. The 20S proteasomes in these cells eventually degrade this disordered region. However, the well-folded ZU5 domain is too large to enter the 20S proteasome’s catalytic chamber, and therefore the 20S proteasome generates the CARD8^p44^ fragment. CARD8^p44^ cannot form an inflammasome but can sequester CT fragments in the DPP8/9 ternary complex, further buffering unstressed cells against inappropriate CARD8 inflammasome activation. In stressed cells (e.g., VbP- or VbP + MeBs-treated cells), however, the ZU5 domain is concomitantly destroyed with the disordered region, leading to the release of the inflammatory CARD8^CT^ fragment. The molecular mechanisms that accelerate the degradation of the ZU5 domain are unknown, but we hypothesize that it either involves the unfolding of the ZU5 domain and/or opening of the 20S proteasome gate (18). On that note, disordered regions dramatically impair the folding of proximal domains (19), and we speculate that CARD8’s disordered region, which is essential for inflammasome activation (8), might play a key role in regulating the folding of the ZU5 domain.

NLRP1 is an inflammasome-forming PRR that is closely related to CARD8 (20). NLRP1 and CARD8 share a similar ZU5-UPA-CARD region, but NLRP1 has N-terminal pyrin (PYD), nucleotide-binding (NACHT), and leucine-rich repeat (LRR) domains instead of a simple disordered sequence. We have proposed that NLRP1 and CARD8 likely both sense the same perturbation in cell homeostasis (8), but that NLRP1, which triggers a more inflammatory response than CARD8 (21), uses its NT domains to further restrain its activation (22). Based on our findings here, we speculate that the NLRP1 NT domains in some way control the rate of NT fragment destruction by the 20S proteasome. The relationship between the 20S proteasome and NLRP1 warrants further investigation.

Overall, the primordial purpose of the CARD8 inflammasome has not been definitively established. The core 20S proteasome rapidly destroys misfolded and disordered proteins, but spares well-folded proteins, and thereby plays a critical role in alleviating proteotoxic stress. Our findings here that the 20S proteasome controls CARD8 suggests that this inflammasome sensor monitors its own ability to fold and avoid destruction. Future studies are needed to determine why this relationship between protein folding and destruction is so closely guarded by the innate immune system.

## Acknowledgements

This work was supported by the NIH (R01 AI137168 and R01 AI163170 to D.A.B.; NIAID K99/R00 AI148598-01 to C.Y.T.; T32 GM007739-Andersen and F30 CA243444 to A.R.G.; NIH T32 GM115327-Tan to E.L.O.-H.; the MSKCC Core Grant P30 CA008748), Gabrielle’s Angel Foundation (D.A.B.), Mr. William H and Mrs. Alice Goodwin, the Commonwealth Foundation for Cancer Research, and The Center for Experimental Therapeutics of Memorial Sloan Kettering Cancer Center (D.A.B.), and the Emerson Collective (D.A.B.). We thank the UC Davis Molecular Structure Facility for the Edman sequencing analysis.

## Author Contributions

D.A.B. conceived and directed the project. J.C.H., A.R.N., A.J.C., A.R.G., C.Y.T., H.C.H., E.L.O.-H., D.P.B., Q.W., and G.H. performed cloning, gene editing, biochemistry, compound screening, and cell biology experiments. D.A.B. and J.C.H. wrote the manuscript.

## Materials and Methods

### Antibodies and reagents

Antibodies used include: GSDMD rabbit polyclonal Ab (Novus Biologicals, NBP2-33422), Cleaved-GSDMD^NT^ rabbit monoclonal Ab (Abcam, Ab215203), PARP rabbit polyclonal Ab (Cell Signaling Tech, 9542), GAPDH rabbit monoclonal Ab (Cell Signaling Tech, 14C10), DPP9 rabbit polyclonal (Abcam, ab42080), HA rabbit monoclonal Ab (Cell Signaling Tech, C29F4), HA mouse monoclonal Ab (Cell Signaling Tech, 6E2), CARD8^CT^ rabbit polyclonal Ab (Abcam, ab24186), CARD8^NT^ rabbit polyclonal Ab (Abcam, ab194585), GFP rabbit polyclonal Ab (Cell Signaling Tech, 2555), V5 mouse monoclonal Ab (Sigma Aldrich, V8012), FLAG mouse monoclonal Ab (Sigma Aldrich, F1804), 20S Proteasome β5 mouse monoclonal (Santa Cruz Biotech, sc-393931), Ubiquitin rabbit polyclonal (Cell Signaling Tech, 3933), IRDye 800CW donkey anti-rabbit (LICOR, 925-32211), IRDye 680RD donkey anti-rabbit (925-68073), IRDye 800CW donkey anti-mouse (925-32212), IRDye 680RD donkey anti-mouse (925-68072). Reagents for immunoprecipitation experiments include anti-FLAG M2 agarose resin (Millipore, A2220), Pierce anti-HA agarose resin (Thermo Scientific, 26181). Other reagents used include Val-boroPro (VbP; Tocris 3719), bortezomib (BORT; Millipore-Sigma, 504314), MG-132 (Sigma-Aldrich, 474787), bestatin methyl ester (MeBs; Sigma, 200485), MLN4924 (Cayman, 15217), dTAG13 (Tocris, 6605), VX-765 (Cayman, 28825), Doxycycline hyclate (Cayman, 14422), FuGENE HD (Promega, E2311), HALT protease inhibitor cocktail (Thermo Scientific, 78430). Human erythrocyte-derived, purified 20S proteasomes used are from South Bay Bio (SBB-PP0005), Aviva Systems Biology (OPED00120), Enzo Life Sciences (BML-PW8720-0050), and R&D Systems (E-360).

### Cell Culture

HEK293T and THP-1 cells were purchased from ATCC. MV4;11 and OCI-AML2 cells were purchased from DSMZ. HEK 293T cells were grown in Dulbecco’s Modified Eagle’s Medium (DMEM) with L-glutamine and 10% fetal bovine serum (FBS). THP-1, MV4;11, and OCI-AML2 cells were grown in Roswell Park Memorial Institute (RPMI) medium 1640 with L-glutamine and 10% fetal bovine serum (FBS). All cells were grown at 37 °C in a 5% CO_2_ atmosphere incubator. Cell lines were regularly tested for mycoplasma using the MycoAlert™ Mycoplasma Detection Kit (Lonza).

### Cloning

Plasmids for full-length and truncated CARD8 were cloned as described previously (8). For constitutive expressions, indicated CARD8 variants were shuttled into pLEX307 vectors that have been modified to contain different N-terminal (e.g., GFP or V5-GFP) or C-terminal (e.g., FLAG) tags using Gateway technology. For generation of CARD8 truncations (e.g., CARD8^ZUC^, CARD8^Δ1-149^), PCR was conducted with primers (with a beginning methionine) that anneal to regions within CARD8. Point mutations (e.g., CARD8 S297A, E274R) were generated using the QuikChange II site-direct mutagenesis kit (Agilent, 200523) following the manufacturer’s protocol. For generation dTAG-CARD8^FL^-V5, the Gateway-compatible pLEX305 N-dTAG was used as the vector backbone to shuttle in CARD8^FL^ that has been attached with a GGGGS-linker sequence followed by a V5-tag and 2 stop codons. The DNA of the MTMR1^M1-Q94^-CARD8^ZUC^ construct was generated by Genscript and subsequently shuttled into the pLEX307 vector. For the generation of CARD8^M1-F161^-GFP, an assembly PCR was performed to fuse CARD8’s disorder to GFP. For the generation of tetracycline (tet)-on-inducible constructs, the Gateway-compatible pINDUCER20 plasmid was used. The DNA of CARD8 lysine mutants (e.g., CARD8 K10R and K26R) were generated by Genscript and shuttled into the indicated vectors (e.g., pINDUCER20 or pLEX307). The CARD8 K10R construct have all 10 lysines N-terminal to the CARD8 autoproteolysis site mutated to arginines (residues 3, 4, 9, 32, 41, 55, 147, 157, 175, 272); and for the CARD8 K26R construct, the rest of the lysines (residues 331, 345, 383, 387, 390, 404, 411, 421, 433, 455, 469, 486, 493, 498, 508, 509). Importantly, every experiment that involve CARD8 lysine mutants and their controls contain a C-terminal GGGGS linker sequence, followed by an HA-tag sequence and 2 stop codons prior to the Gateway *attB2* recombination site. Therefore, the C-terminal sequences following CARD8’s open reading frame do not code for any lysines.

### Transient transfections

HEK 293T cells were plated in 6-well culture plates at 5.0 × 10^5^ cells/well in DMEM. The next day, the indicated plasmids were added to a total of 2.0 ug DNA (with pLEX307 RFP as the filler plasmid) in 125 uL Opti-MEM and transfected using FuGENE HD (Promega) according to the manufacturer’s protocol. Unless indicated otherwise, 2 ug of each plasmid construct was used. For experiments involving constitutive plasmid expressions, the cells were incubated for an additional 48 hours before their harvest. For experiments involving tet-inducible plasmid constructs, the cells were treated with DOX at 1 ug/mL (and/or with other compounds) 20 hours after transfection. The cells were incubated for an additional 24 hours before their harvest.

### CRISPR/Cas9 gene editing

To generate *CARD8* knockouts in THP-1 cells, 1.5 × 10^6^ cells stably expressing Cas9 (3) were infected with lentivirus containing sgRNA plasmids (packaged in HEK 293T cells using Fugene HD and 2 ug of the vector, 2 ug psPAX2, and 1 ug pMD2.G). After 48h, cells were selected with hygromycin (100 ug/mL) until control cells died. Single cell clones were isolated by serial dilution and confirmed by Western blotting.

### Generation of stable cell lines

Indicated expression plasmids (e.g., pLEX307 CARD8^WT^ HA) were packaged into lentivirus in HEK 293T cells as described above. It should be noted that all CARD8 plasmids introduced into CARD8^−/−^ THP-1 have silent mutations at residues F211 and S213 (generated by the Quikchange kit). 1.5 × 10^6^ cells of CARD8^−/−^ THP-1 were then infected with the virus, and after 48h, selected with puromycin (0.5 ug/mL) until control cells died.

### LDH cytotoxicity, immunoblotting, and FLAG/HA immunoprecipitation

HEK 293T cells were transiently transfected and compounds treated as indicated. THP-1 cells were plated in 12-well culture plates at 0.5 × 10^6^ cells/well and treated with compounds as indicated. Supernatants were analyzed for LDH levels using the Pierce LDH Cytotoxicity Assay Kit (Life Technologies). LDH levels were quantified relative to a lysis control where cells were lysed in 20 μL of a 9% Triton X-100 solution. For immunoprecipitation experiments, cell pellets were sonicated and centrifuged at 3220 x *g* for 5 min. The clarified lysates were retained and were subsequently incubated with 40 μL of anti-FLAG-M2 agarose resin (Sigma) for 1.5 hours at room temperature or anti-HA agarose resin (Thermo Scientific) for overnight at 4C. After washing with three rounds of 100 uL of PBS, bound proteins were eluted by incubating resin with 100 μL of PBS containing 150 ng/uL 3X-FLAG peptide or 1 mg/mL of HA peptide for 1 hour at room temperature. An equal volume of 2X sample loading dye was added to the eluate and incubated at 95°C for 10 min. For immunoblotting, cells were washed with two rounds of PBS (pH = 7.4), resuspended in PBS that were added with 1X HALT protease inhibitor, lysed by sonication, and briefly clarified by centrifuging at 1000 x *g* for 10 min. Protein concentrations were determined and normalized using the DCA Protein Assay kit (Bio-Rad). The samples were separated by SDS-PAGE, immunoblotted, and visualized using the Odyssey Imaging System (Li-Cor).

### 20S proteasome assays

HEK 293T cells were transiently transfected with the indicated plasmid constructs. After 48 hours, the lysates were incubated with anti-FLAG or anti-HA beads before eluting with the corresponding peptides to enrich for the expressed proteins. The eluent is composed of 20 mM Tris-HCl, 20 mM NaCl, 10 mM MgCl2, 1 mM DTT, pH 7.5. As in Figures 3A, C, D, and E, the eluates were diluted with the same eluent three times before incubating with or without 200 nM of purified 20S proteasomes. As in Fig. 3B and Fig. S3, the eluates were subjected to a size-exclusion filtration step (to rid the eluate of excess FLAG or HA peptide), before quantifying the protein concentration with DCA assay. CARD8 (800 nM) was then incubated with 20S proteasomes (100 nM). All 20S reactions were incubated at 37C for four hours, shaking at 500 RPM before quenching with 2X loading dye prior to immunoblotting analysis.

### Edman degradation

HEK 293T cells were transiently transfected with a plasmid encoding CARD8^FL^ WT FLAG. After 48 hours, the lysates were purified with anti-FLAG beads. Then, the eluate was added with 2X loading dye and boiled, and the protein was separated by SDS-PAGE. The gel was then transferred onto a PVDF membrane and stained with Ponceau S. The bands of interest were excised and sent to the UC Davis Molecular Structure Facility for Edman sequencing analysis.

### Statistical analysis

Student’s two-sided T-tests were performed in Figures 1E,F; 4A,B; S1C; and S4E. *P* values less than 0.05 were significant. Graphs and error bars represent means ± SD of a single experiment representative of three or more independent experiments unless stated otherwise. The investigators were not blinded in all experiments. All statistical analysis was performed using GraphPad Prism 9.

### Data availability

All data in this study are available within the paper, Supplementary information, and/or from the corresponding author on reasonable request.

---

**Figure S1.**
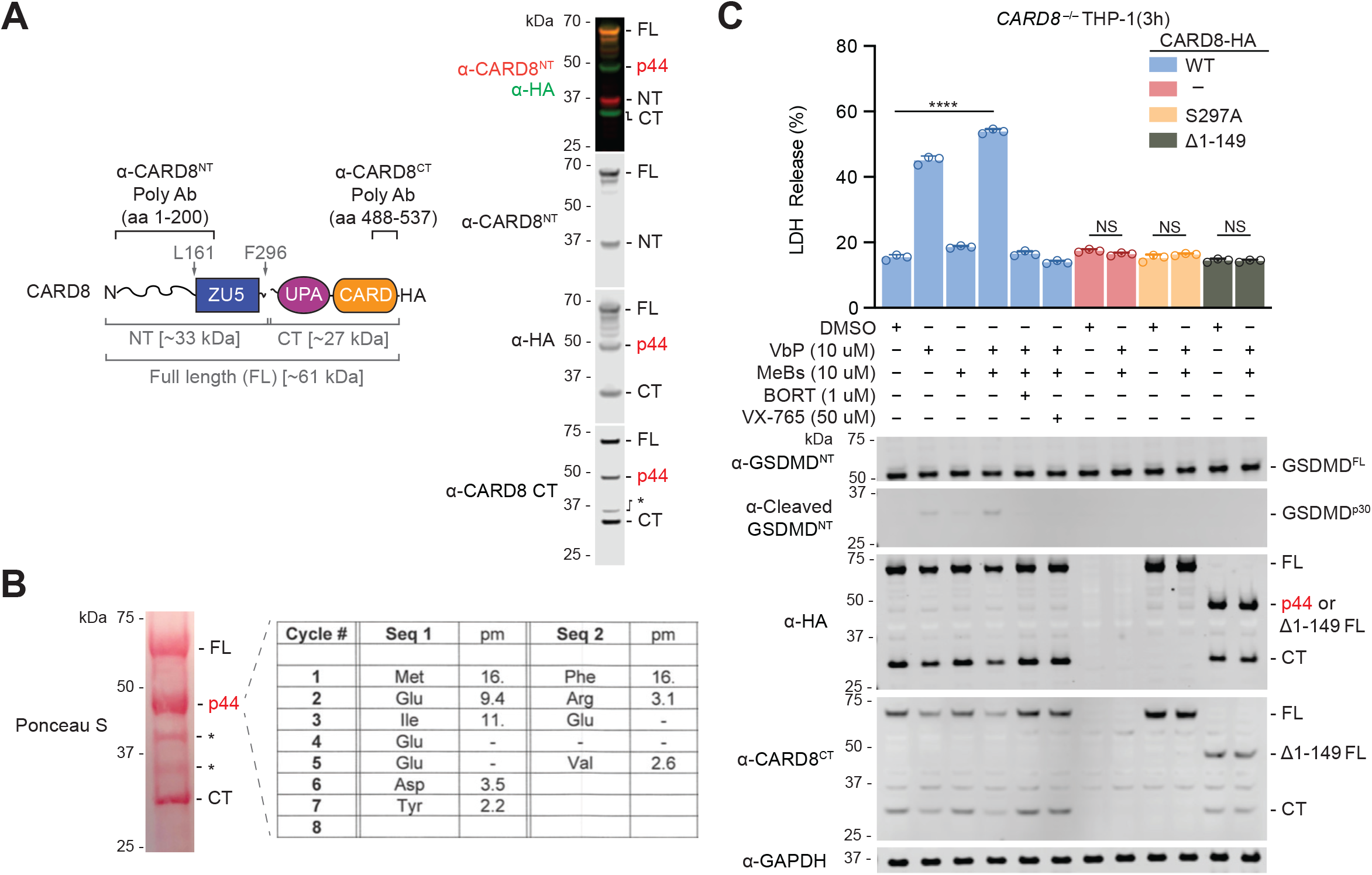
Identification and characterization of CARD8^p44^. (**A**) The specific residues that the antibodies target on CARD8 are depicted on the cartoon (left). HEK 293T cells were transiently transfected with a plasmid encoding C-terminally HA-tagged CARD8^FL^ WT construct. The lysate was analyzed by immunoblotting (right). (**B**) C-terminally FLAG-tagged CARD8^WT^ expressed from HEK 293T cells were purified with anti-FLAG beads. The excised, ponceau-stained band was analyzed by Edman degradation (right). pm denotes picomoles. (**D**) *CARD8*^*−/−*^ THP-1 cells stably expressing CARD8 WT, S297A, or Δ1-149 were treated with indicated compounds for 3 h before LDH release and immunoblotting analyses. **** *p* < 0.0001, by Student’s two-sided *t*-test. NS, not significant. Data are means ± standard deviation (SD) of 3 replicates. All data, including immunoblots, are representative of 3 or more independent experiments.

**Figure S2.**
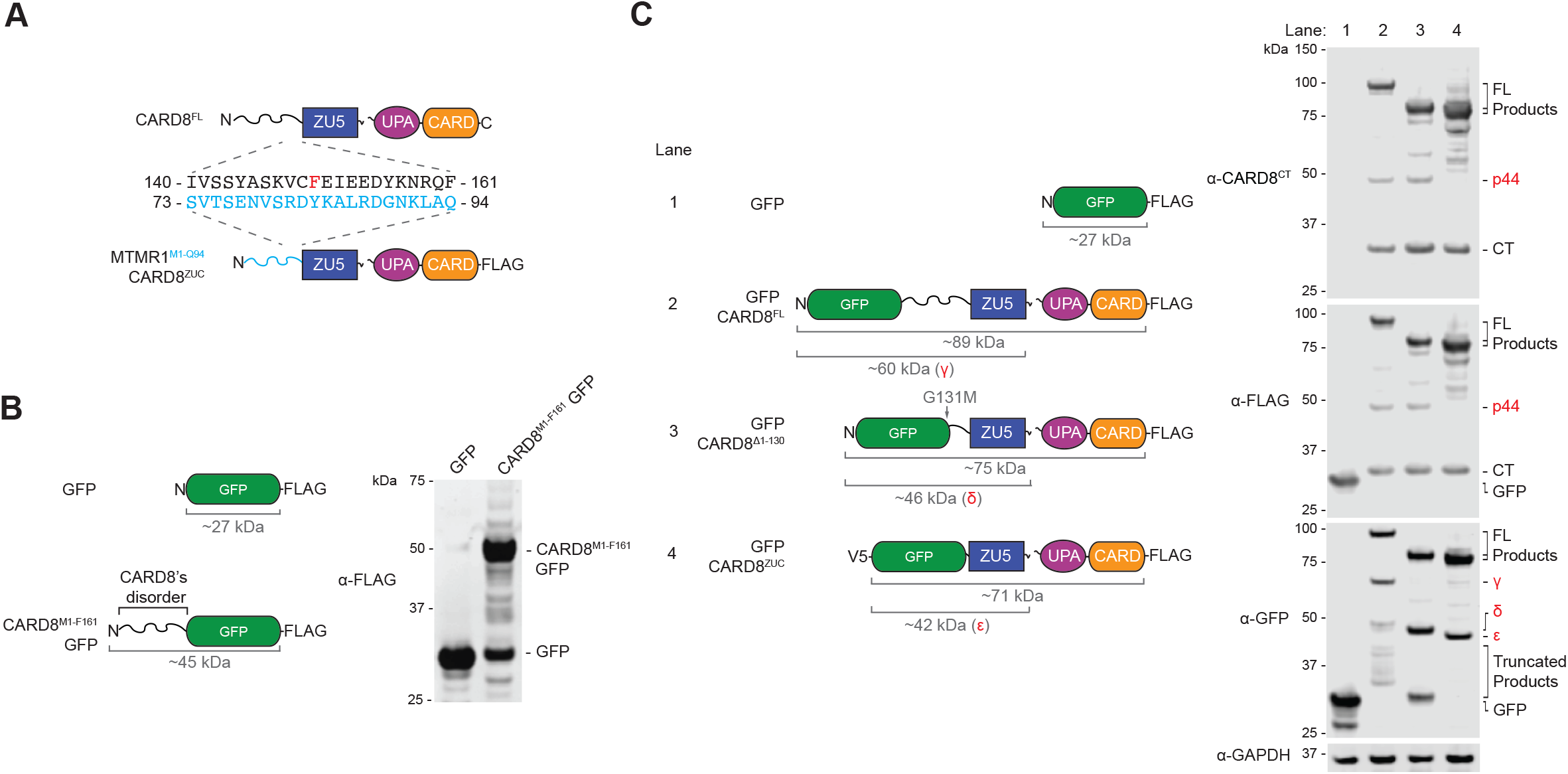
The removal of CARD8’s disordered region is sequence independent. (**A**) Sequence alignment of the twenty-two amino acid residues prior to the ZU5 domain of each indicated construct. (**B and C**) HEK 293T cells were transiently transfected with a plasmid encoding the indicated constructs (left). Lysates were analyzed by immunoblotting (right). All immunoblots are representative of 3 or more independent experiments.

**Figure S3.**
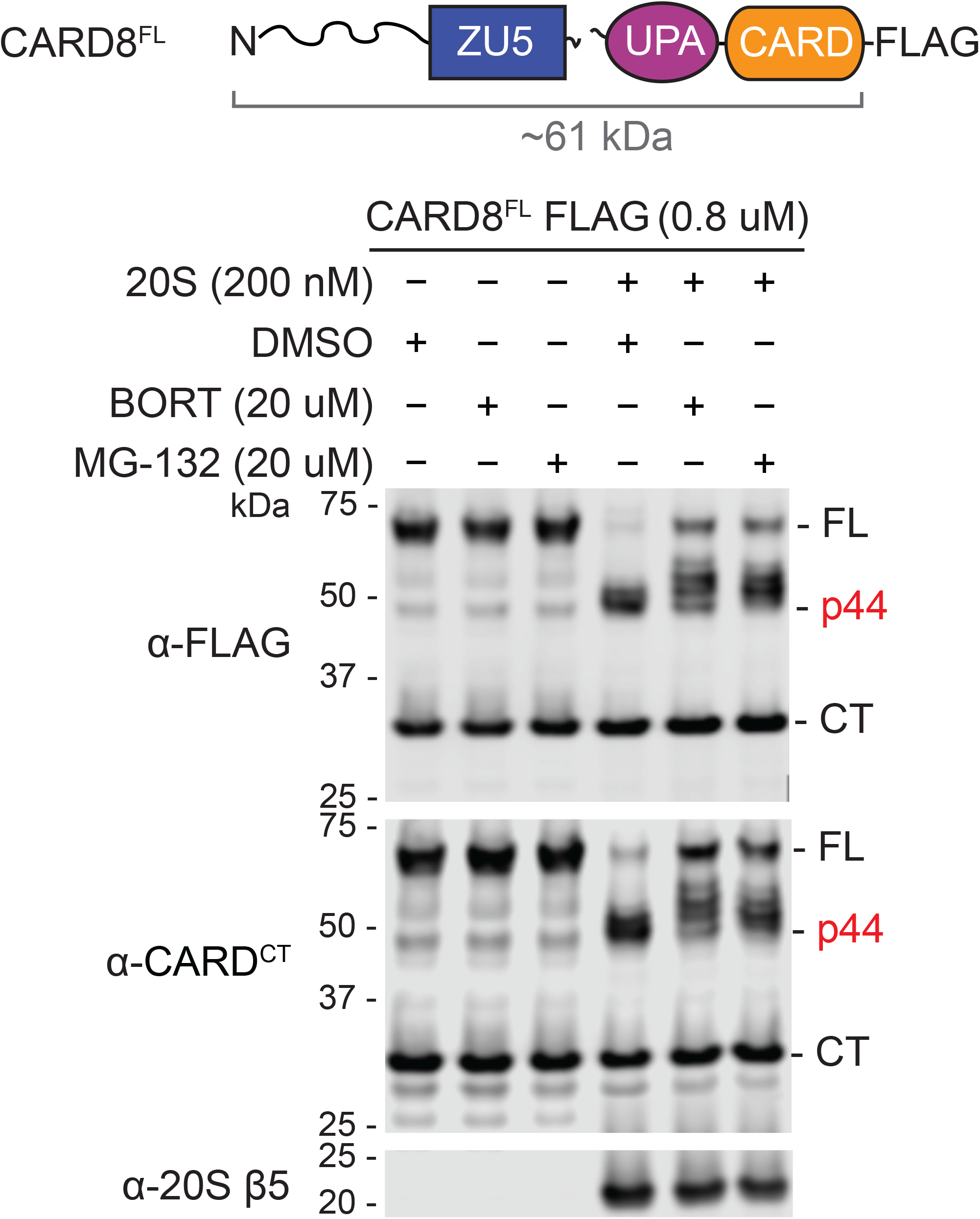
Proteasome inhibitors slow the degradation of CARD8^FL^. Purified CARD8^FL^ WT protein incubated with the indicated inhibitors and purified 20S proteasomes for 4 h. Reactions were quenched with 2X loading dye prior to immunoblotting analysis. The immunoblot is representative of 3 independent experiments.

**Figure S4.**
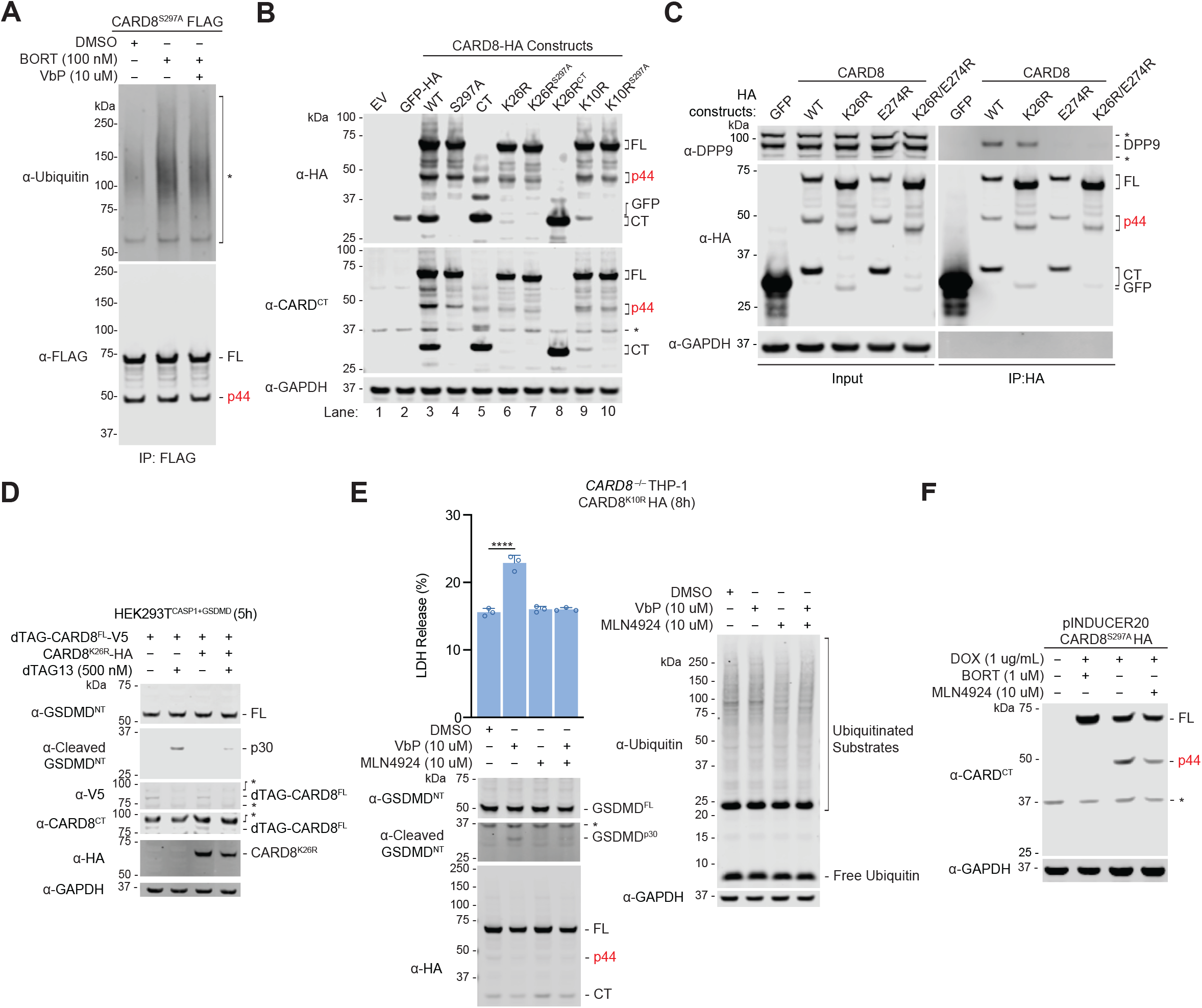
Characterization of CARD8 lysine mutants. (**A**) HEK 293T cells were transiently transfected with C-terminally FLAG-tagged CARD8^FL^ S297A and incubated for 24 h. In the last 6 hours, the cells were treated as indicated before lysates were harvested. FLAG-tagged proteins were then enriched on anti-FLAG beads, and the eluates were analyzed by immunoblotting. It should be noted that the signals detected on the anti-ubiquitin immunoblots in the BORT-treated samples were likely due to ubiquitinated substrates that bound non-specifically to the anti-FLAG beads, as no laddering pattern of CARD8 (in both the anti-UBQ and FLAG immunoblots) was observed. (**B**) HEK 293T cells were transiently transfected with the indicated plasmid constructs prior to immunoblotting analyses. It should be noted that the laddering of the CT protein is likely due to incomplete dissociation of its oligomers by the reducing agent and boiling procedure. (**C**) HEK 293T cells were transiently transfected with the indicated constructs. Lysates were harvested and HA-tagged proteins were enriched. The input and elution fractions were analyzed by immunoblotting. The CARD8 E274R mutation renders its inability to bind to DPP8/9. (**D**) HEK 293T^CASP1+GSDMD^ cells were transfected with plasmids encoding the indicated plasmids. After 24 h, cells were treated with dTAG-13 for 5 h before immunoblotting analyses. (**E**) *CARD8*^*−/−*^ THP-1 cells stably expressing CARD8^FL^ K10R were treated with the indicated compounds prior to LDH release and immunoblotting analyses. (**F**) A tet-on plasmid encoding CARD8 was transfected into HEK 293T cells. After 20 h, cells were co-treated with the indicated drugs for 24 h before lysates were analyzed by immunoblotting. All immunoblots are representative of 3 or more independent experiments.

## Notes

### Competing Interest Statement

The authors have declared no competing interest.

